# Depicting pseudotime-lagged causality across single-cell trajectories for accurate gene-regulatory inference

**DOI:** 10.1101/2022.04.25.489377

**Authors:** Caleb C. Reagor, Nicolas Velez-Angel, A. J. Hudspeth

**Affiliations:** Howard Hughes Medical Institute and Laboratory of Sensory Neuroscience, The Rockefeller University, New York, NY, USA; Tri-Institutional PhD Program in Computational Biology and Medicine, New York, NY, USA

## Abstract

Identifying the causal interactions in gene-regulatory networks requires an accurate understanding of the time-lagged relationships between transcription factors and their target genes. Here we describe DELAY, a convolutional neural network for the inference of gene-regulatory relationships across pseudotime-ordered single-cell trajectories. We show that combining supervised deep learning with joint-probability matrices of pseudotime-lagged trajectories allows the network to overcome important limitations of ordinary Granger causality-based methods, such as the inability to infer cyclic relationships such as feedback loops. Our network outperforms several common methods for inferring gene regulation and predicts novel regulatory networks from scRNA-seq and scATAC-seq datasets given partial ground-truth labels. To validate this approach, we used DELAY to identify important genes and modules in the regulatory network of auditory hair cells, as well as likely DNA-binding partners for two hair cell cofactors (Hist1h1c and Ccnd1) and a novel binding sequence for the hair cell-specific transcription factor Fiz1. We provide an open-source implementation of DELAY at https://github.com/calebclayreagor/DELAY.

## Introduction

Single-cell sequencing technologies can provide detailed data for the investigation of heterogeneous populations of cells collected at specific times—so-called “snapshots”—during cellular differentiation or dynamic responses to stimulation^1^. However, owing to inherent delays in molecular processes such as transcription and translation, static measurements from individual cells cannot reveal the causal interactions governing cells’ dynamic responses to developmental and environmental cues^2–4^. Because population-level heterogeneity in tissues often reflects the asynchronous progression of single cells through time-dependent processes, observed patterns of gene expression can nonetheless indicate the stages of development to which individual cells belong^5^. Many algorithms exploit these cell-to-cell differences to infer dynamic trajectories and reconstruct cells’ approximate temporal progressions along inferred lineages in pseudotime^6, 7^.

Several methods for gene-regulatory inference rely on pseudotime in Granger causality tests, which try to determine whether new time series can add predictive power to inferred models of gene regulation^8, 9^. However, Granger causality-based methods can be error-prone when genes display nonlinear or cyclic interactions, or when the sampling rate is uneven or too low^9–12^. Because pseudotime trajectories exhibit these problems, Granger causality-based methods often underperform model-free approaches that exploit pure statistical dependencies in gene-expression data^9, 13, 14^.

By contrast, deep learning-based methods make no assumptions about the temporal relationships or connectivity between genes in complex regulatory networks; instead, these data-driven approaches learn general features of regulatory interactions^15, 16^. Here we describe a deep learning-based method termed DELAY (*<u>De</u>*picting *<u>La</u>*gged Causalit*<u>y</u>*) that learns gene-regulatory interactions from discrete joint-probability matrices of paired, pseudotime-lagged gene-expression trajectories. Our data suggest that DELAY can address many shortcomings of current Granger causality-based methods and provide a useful, complementary approach to overcome common limitations in the inference of gene-regulatory networks from single-cell data.

## Results

### A convolutional neural network predicts pseudotime-lagged gene-regulatory relationships

To predict gene-regulatory relationships from single-cell data, we developed a convolutional neural network based on Granger causality^17^. The input to DELAY consisted of stacks of two-dimensional joint-probability matrices for pairs of transcription factors, putative target genes, and highly correlated “neighbor” genes^18^. We constructed the input matrices by aligning the gene-expression trajectories of a transcription factor *A* at several lagged positions in pseudotime relative to a target gene *B* to generate joint-probability distributions from two-dimensional histograms of gene coexpression at each lag (Fig. 1a). Each input matrix consisted of the L2-normalized cell-number counts from a two-dimensional gene-coexpression histogram with 32 fixed-width bins in each dimension, spanning each gene’s minimum and maximum expression values. Although the marginal probability distributions for both *A* and *B* remained essentially unchanged at each lag except for cells lost at the leading and lagging edges of the shifted trajectories, realigning the gene expression in pseudotime altered key features of the resulting joint-probability matrices. In other words, causally related genes share important pseudotime-lagged patterns of gene coexpression with nearby cells in single-cell trajectories.

**Figure 1:**
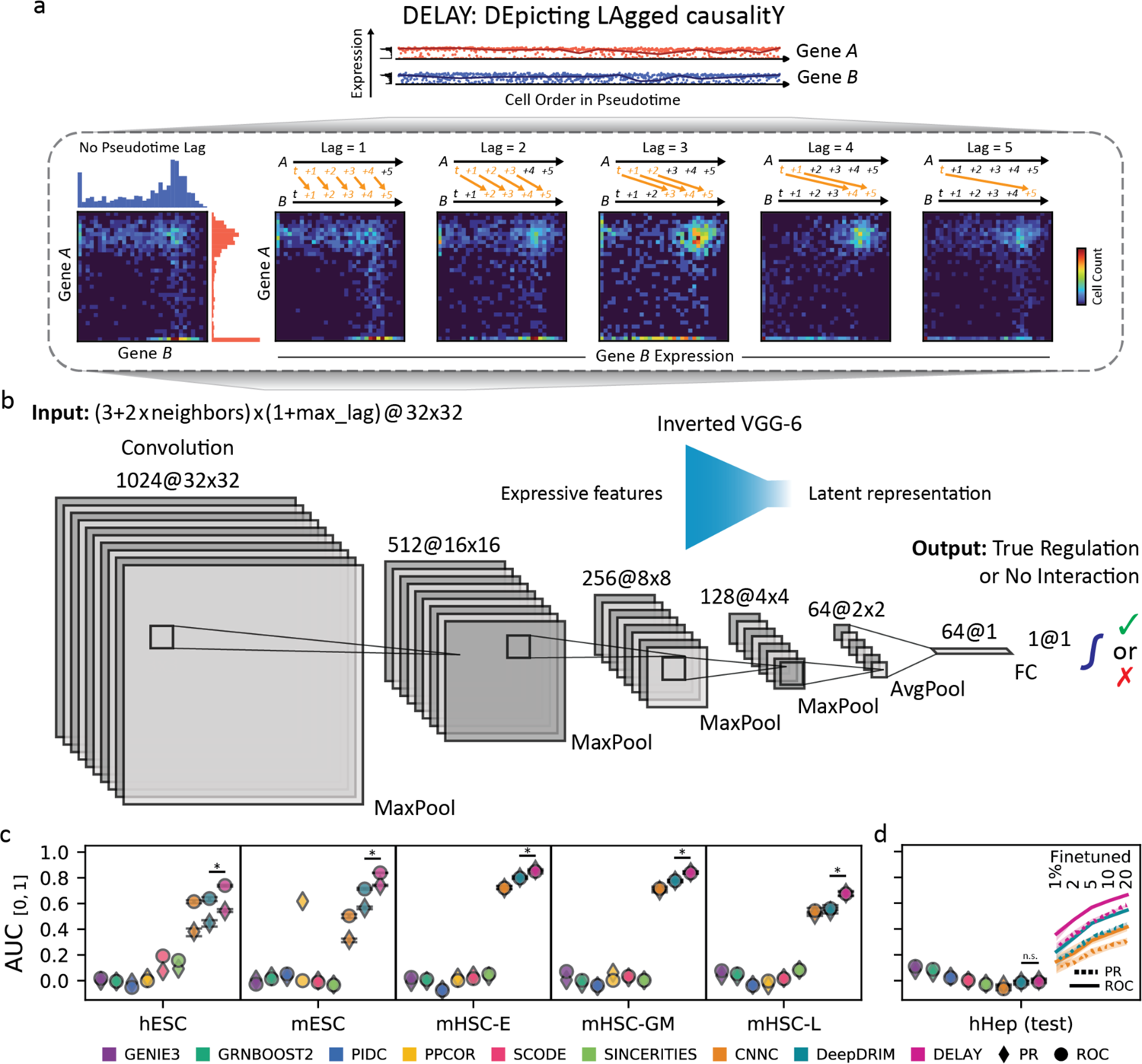
Pseudotime-lagged causality allows accurate inference of gene-regulatory networks. **a.** Shifting the gene-expression trajectory of transcription factor *A* forward in pseudotime with respect to that of target gene *B* generates a series of unique joint-probability distributions. The resulting joint-probability matrices contain gene-coexpression signatures—in the upper-left, upper-right, and lower-right regions—that indicate *A* directly regulates *B*. **b.** The inverted architecture of DELAY is wide at the beginning of the network and progressively narrows to a latent space followed by a linear classifier and sigmoid activation function to generate gene-regulation probabilities. The network uses leaky ReLU activations and padded 3×3 convolutions throughout. **c.** When trained on five benchmark scRNA-seq datasets, DELAY outperforms eight of the most popular methods for inferring gene-regulatory interactions. **d.** DELAY does not immediately generalize to a new testing dataset but outperforms all other methods if fine-tuned on a small percentage of the new data. The error bars and markers in **c** and **d** show the full range and average performance for each neural network across five cross-validated models. The values for areas under curves are normalized between a random predictor and perfect performance. The statistical significance between DELAY and the next-best neural network was assessed using a one-sided Wilcoxon signed-rank test (*, *P*≤0.05). PR, precision-recall; ROC, receiver operating characteristic.

Using ground-truth labels from cell type-specific chromatin-immunoprecipitation sequencing (ChIP-seq)^14^, we conducted supervised learning to train our neural network to predict whether *A* directly regulates *B*. This procedure resembled a regression in a Granger causality test^17^, in which values of a time series *Y* at timepoints *y*_t_ are regressed against values from another time series *X* at timepoints *x_t_*, *x_t-1_*, *x_t-2_*, …, *x_t-T_* up to some maximum lag *T* to determine whether any time-lagged values of *X* add explanatory power to *Y*’s autoregressive model. DELAY likewise learned higher weights for gene-coexpression matrices at specific pseudotime lags that indicated the true regulatory relationship between genes. After comparing several neural network architectures, we selected a six-layered convolutional network trained on pseudotime-aligned (*T*=0) and five pseudotime-lagged (*T*∈{1,2,3,4,5}) gene-coexpression matrices to predict direct gene-regulatory relationships (Fig. 1b, Supplementary Fig. 1). We also trained the network on lagged-coexpression matrices of two highly correlated neighbor genes per gene, that is, the two transcription factors with the highest cross-correlation with *A* and *B* along the single-cell trajectory. Because the neighbor genes can present stronger alternatives to the primary hypothesis that *A* directly regulates *B*, including these matrices for *A* and *B versus* their highly-correlated transcription factors reduced false-positive predictions.

### DELAY outperforms several common methods of gene-regulatory inference

We trained DELAY on gene-expression datasets from human embryonic stem cells (hESCs)^19^, mouse embryonic stem cells (mESCs)^20^, and three lineages of mouse hematopoietic stem cells (mHSCs)^21^. We generated separate training datasets for each of the hematopoietic lineages, and all datasets contained at least four hundred cells per lineage (Table 1). Each trajectory was oriented according to known experimental timepoints or precursor-cell types and lineages, and pseudotime values were inferred separately with Slingshot^6^ for each lineage (Supplementary Fig. 2). We chose these datasets because the cell types and trajectories are well characterized and offer cell type-specific ChIP-seq data to generate ground-truth networks^14^. Although the three hematopoietic datasets contained similar numbers of examples of true regulation and no interaction, both of the embryonic datasets were class-imbalanced and contained fewer examples of true regulation. To avoid overfitting, we performed five-fold cross validation on randomly segregated 70 %-30 % splits of all possible gene-pair examples, that is, both true regulation and no interaction, from each dataset and trained models on the combined 70 % splits across all five datasets.

**Table 1:** Summary of the datasets used to train, test, and evaluate the neural network. Columns to the left (Cell Type to Cells) describe the single-cell datasets used to train, test, and evaluate DELAY, and columns to the right (TFs to Density) describe the resulting gene-regulatory networks, given the selected transcription factors and target genes. For three of the networks, differentially-expressed transcription factors without target genes (in parentheses) were included in our analysis as target genes, but not as transcription factors. All transcription factors were also included as target genes, and gene-pair examples included all transcription factor-target gene pairs, including examples of both true regulation (Edges, from ChIP-seq ground-truth data) and no interaction. Gene Expression Omnibus (GEO) accession numbers are listed for each dataset. Where indicated (*), network descriptions refer to partial ground-truth networks. hESC, human embryonic stem cell; mESC, mouse embryonic stem cell; mHSC, mouse hematopoietic stem cell; E, erythroid lineage; GM, granulocyte-monocyte lineage; L, lymphoid lineage; hHep, human hepatocyte; mOHC, mouse outer hair cell; TC, time course; S, snapshot; TF, transcription factor.

| Datasets |  |  |  |  |  | Networks |  |  |  |
| --- | --- | --- | --- | --- | --- | --- | --- | --- | --- |
| Cell Type | TC/S | GEO | Set | Cells | TFs | Targets | Examples | Edges | Density |
| hESC | TC | GSE75748 | Train | 758 | 37 (4) | 541 | 20,017 | 3,166 | 15.82% |
| mESC | TC | GSE98664 | Train | 421 | 125 (17) | 642 | 80,250 | 20,580 | 25.64% |
| mHSC<br>-E | S | GSE81682 | Train | 1,071 | 33 | 533 | 17,589 | 9,592 | 54.53% |
| -GM | S | GSE81682 | Train | 889 | 23 | 523 | 12,029 | 6,587 | 54.76% |
| -L | S | GSE81682 | Train | 847 | 16 | 516 | 8,256 | 4,026 | 48.76% |
| hHep | TC | GSE81252 | Test | 425 | 33 (3) | 536 | 17,688 | 6,171 | 34.89% |
| mOHC<br>-scRNA | TC | GSE137299 | Eval | 1,563 | 56 | 556 | 1,112* | 166* | 14.93%* |
| -scATAC | S | GSE157398 | Eval | 391 | 48 | 382 | 764* | 157* | 20.55%* |

Our network outperformed eight of the most popular approaches for inferring gene-regulatory relationships, including two deep convolutional neural networks^18, 22^ and six other methods^8, 23–27^ (Fig. 1c). We measured the performance of the six methods across the combined training and validation splits for each dataset, then compared the results with the cross-validated performance of the deep learning-based methods on the held-out examples alone. With one exception, the deep learning-based methods outperformed all others according to the areas under both the precision-recall (PR) and receiver operating characteristic (ROC) curves. Moreover, DELAY outperformed all the other methods according to both metrics but performed slightly worse by area under the PR curve than by area under the ROC curve for the class-imbalanced embryonic datasets. Even though one of the deep learning-based methods, DeepDRIM, was trained on fivefold as many neighbor-gene matrices as DELAY, it still performed second- or third-best behind DELAY for all metrics. Together, these results suggest that DELAY outperforms the eight other methods because it learns important features from pseudotime-lagged gene-coexpression matrices.

### Transfer learning allows DELAY to predict novel gene-regulatory networks from new single-cell datasets

To test whether DELAY generalizes to new datasets, we examined the human hepatocyte (hHep) gene-regulatory network using an additional dataset with over four hundred single cells and known ground-truth interactions from ChIP-seq data^14, 28^. We inferred the network by the previous methods and found that tree- and mutual information-based methods performed slightly better than deep learning-based methods, which performed comparably to random predictors (Fig. 1d). To determine whether this lack of generalizability arises from batch effects in the single-cell data, we used transfer learning to fine-tune the three deep learning-based methods on small fractions of the new dataset. After training each network on only 1 % of the new examples, their performance matched or exceeded that of the other methods, and training on up to 20 % of the examples yielded further performance increases. As observed previously, DELAY outperformed every other deep learning-based method. We did not observe any overfitting in the networks’ fine-tuning curves for the mean validation metric, suggesting that the networks recognized general features of direct gene regulation associated with strong, well-optimized minima in the training loss. Moreover, these results demonstrate that DELAY can accurately predict gene-regulatory relationships from new datasets with partially known ground-truth labels.

### DELAY performs well with new input configurations, augmented matrices, and modified datasets

Using different numbers of pseudotime-lagged gene-coexpression matrices or neighbor-gene matrices, as well as examples from modified datasets with fewer cells or additional gene dropouts, we next examined the performance of DELAY across various input configurations. We employed the original datasets to train new cross-validated models on gene-coexpression matrices of up to ten pseudotime lags (Fig. 2a) or up to ten neighbor genes (Supplementary Fig. 3), as well as on gene-coexpression matrices with varying dimensions and resolutions. Training DELAY on at least one pseudotime-lagged matrix or with at least one neighbor gene greatly increased the network’s performance across all datasets. Although training DELAY on up to ten pseudotime-lagged matrices resulted in comparable or slightly better performance across all datasets, training the network on more than two neighbor-gene matrices per gene decreased performance in some instances. Adding channel masks for specific lagged matrices suggested that DELAY relies on redundancies across all available lagged inputs (Supplementary Fig. 4).

**Figure 2:**
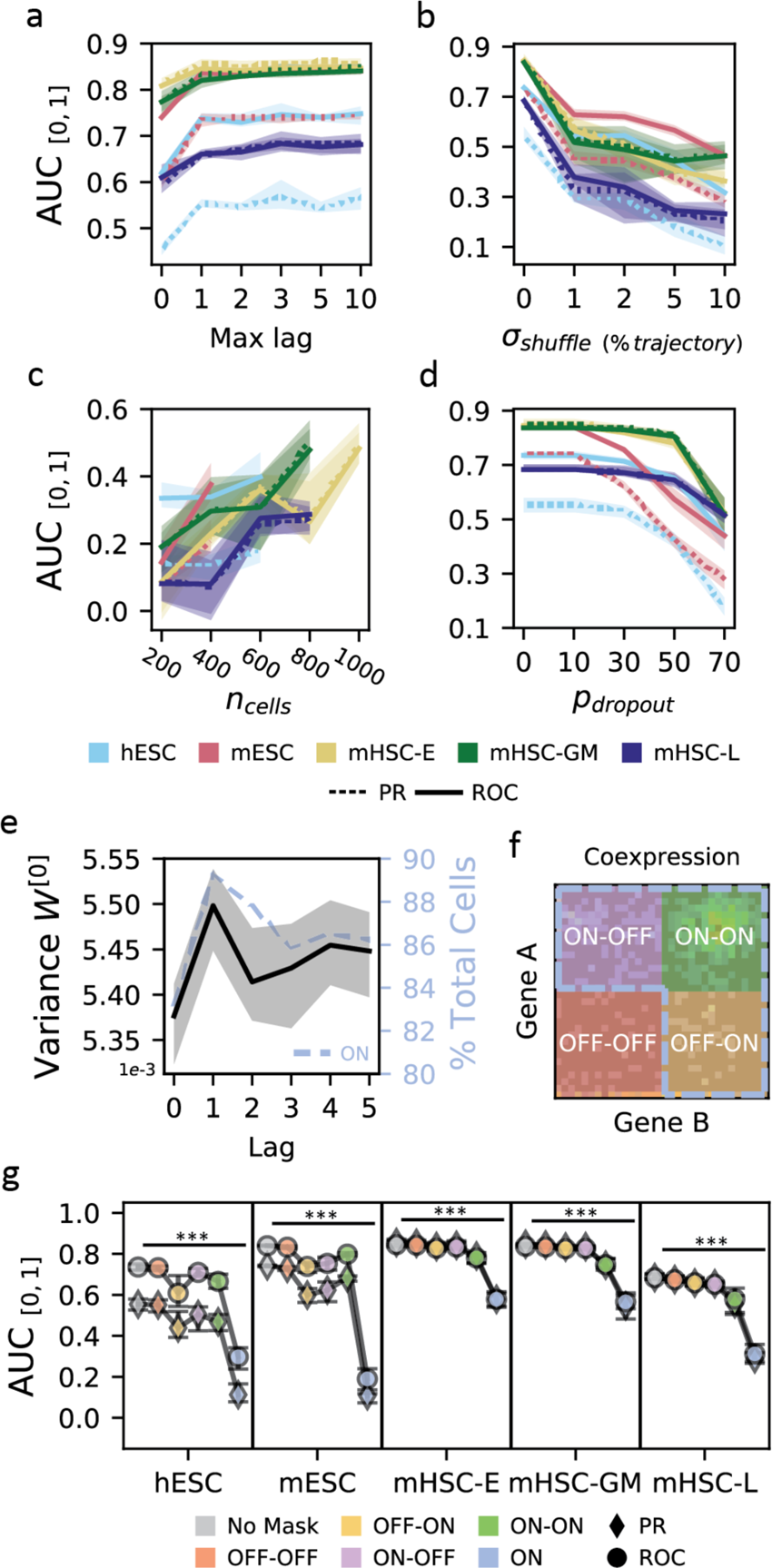
The neural network relies on cell order in pseudotime and gene-expression strength. **a.** Training new models of DELAY on increasing numbers of pseudotime-lagged matrices gives the largest performance increase when using up to a single lag. **b.** Reordering single cells in pseudotime sharply decreases performance, suggesting that DELAY relies on the specific ordering of adjacent cells in each trajectory. **c, d.** The network is sensitive to random down-sampling of cells across datasets (**c**) but relatively more robust to induced, additional gene dropouts in weakly-expressing cells (**d**), suggesting DELAY relies heavily on highly-expressing cells. **e, f.** The network learns larger input weights for lagged matrices of the first pseudotime lag (**e**), which also contain on average more cells in the combined “ON” region (A_ON_∩ B_ON_; blue dotted outline, **f**) across training datasets (blue dotted line, **e**). The combined “ON” region is comprised of the upper-left “ON-OFF” quadrant (A_ON_∩ B_OFF_; purple, **f**), upper-right “ON-ON” quadrant (A_ON_∩ B_ON_; green, **f**), and lower-right “OFF-ON” quadrant (A_OFF_∩ B_ON_; yellow, **f**). **g.** Masking different regions of the input matrices shows that the network relies heavily on the combined “ON” region. Error bars and markers in **a-e** and **g** show the full range and average values across five cross-validated models. The statistical significance in **g** was assessed with a Kruskal-Wallis test (***, *P*≤0.001).

We also characterized DELAY’s performance on pseudotime-shuffled trajectories and observed a sharp decrease in performance after reordering nearby cells in each trajectory (Fig. 2b). This result suggests that the network relies heavily on the specific ordering of adjacent cells in each trajectory. Upon examining DELAY’s performance on modified datasets containing fewer single cells (Fig. 2c) or additional gene-dropout noise (Fig. 2d), we also discovered that the network was more sensitive to down-sampling of single cells than to gene-expression losses in low-expressing cells alone. These results indicate that DELAY relies more heavily on highly expressing cells and therefore learns larger input weights for the first lagged input because on average that input contains stronger features of gene activation than other lags (Figs. 2e,f). To investigate this hypothesis, we used augmented input matrices with masked regions to show that DELAY relies heavily on the combined upper-left, upper-right, and lower-right regions (A_ON_ ∪ B_ON_) across all lagged matrices (Fig. 2g). By performing a *post hoc* analysis of the statistical dispersion across correctly inferred gene-pair examples, we additionally found that DELAY performs best on transcription factors with stable gene expression along single-cell trajectories (Supplementary Fig. 5).

### DELAY recognizes causal relationships in higher-order and cyclic gene-regulatory interactions

To further investigate how the selection of neighbor genes can alter DELAY’s internal representations and subsequent gene-regulatory inferences, we modified gene-pair examples to exclude all neighbor genes with known higher-order interactions across several classes of three-gene motifs. First, we used DELAY to infer gene regulation across several classes of potential higher-order interactions (Fig. 3a), including mutual interactions (MIs), feedback loops (FBLs), and three classes of feedforward loops (FFLs). Unlike ordinary Granger causality, DELAY performs well across cyclic interactions including 2-cycles (MIs) and 3-cycles (FBLs), which are ubiquitous among gene-regulatory networks^29^. We next excluded all input matrices for neighbor genes involved in potential FBLs and FFLs to ascertain how DELAY performs in their absence. Because we saw a significant decrease in performance upon excluding these neighbors in most cases of shared and sequential regulation but not downstream targets, DELAY apparently recognizes the differences between input matrices of causally related genes and those of merely correlated genes. Moreover, these results suggest that although our network uses principles of Granger causality, it can achieve true causal inference for genes involved in several classes of higher-order and cyclic regulatory interactions by avoiding several limiting assumptions of ordinary Granger causality.

**Figure 3:**
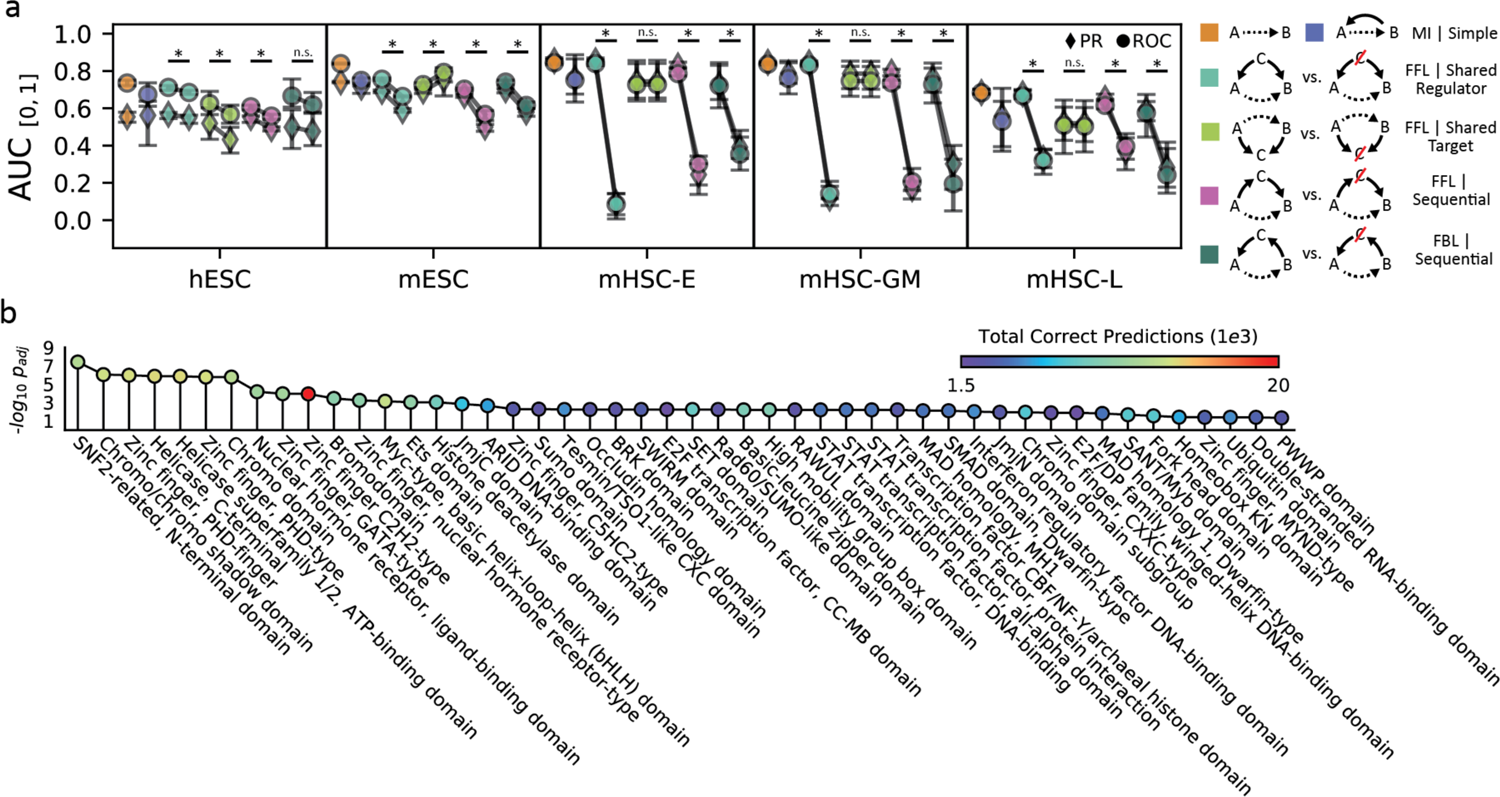
DELAY uncovers causal, cyclic, and context-specific gene-regulatory relationships. **a.** The performance of the network on putative pairs of transcription factors and target genes (dashed lines, legend) in two- and three-gene motifs with additional, known regulatory relationships (solid lines, legend) suggests that DELAY distinguishes between various types of higher-order and cyclic gene regulation. Upon exclusion of neighbor gene *C* from the input features, performance declines significantly in cases of shared and sequential regulation (cyan and pink/teal, respectively) but not for shared downstream targets (lime). **b.** Gene ontology-term enrichment indicates that DELAY performs best on zinc-finger proteins (PHD-type, GATA-type, C2H2-type, NHR-type, C5HC2-type), bHLH factors (Myc-type), and other chromatin remodelers (SNF2-related, chromo domain, bromodomain, HDA domain, helicase). Error bars and markers in **a** show the full range and average performance across five cross-validated models. The statistical significance in **a** was assessed with a one-sided Wilcoxon signed-rank test (*, *P*≤0.05) and in **b** with a Fisher’s exact test (*P*_adjusted_<0.05). MI, mutual interaction; FFL, feedforward loop; FBL, feedback loop.

### DELAY performs best on zinc-finger proteins, bHLH factors, and other chromatin remodelers

To explore how DELAY performs on classes of transcription factors containing different types of DNA-binding domains, we analyzed the enrichment of gene-ontology (GO) terms across correctly inferred transcription factors (Fig. 3b). We found that DELAY performs best on zinc-finger proteins, bHLH factors, and other chromatin remodelers. Although DELAY performs well on C2H2-type zinc-finger proteins in terms of the total number of correct predictions, we found that it performs better on other chromatin remodelers and plant homeodomain (PHD) zinc-finger proteins by the overall term enrichment. Interestingly, these results indicate that training the network on cell type-specific ChIP-seq data allows the network to identify regulatory relationships involving some non-sequence-specific transcription factors and cofactors that nevertheless associate with specific targets at preferred chromatin conformations (Supplementary Table 1).

### Predicting the gene-regulatory network of auditory hair cells through multi-omic transfer learning

We next sought to generate predictions for a cell type with complex but incompletely characterized gene-regulatory dynamics. We devised a pipeline for multi-omic transfer learning to infer the gene-regulatory network for developing mouse outer hair cells (mOHCs), the mechanical amplifiers of the inner ear. We used both gene-expression^30^ and chromatin-accessibility^31^ datasets to fine-tune our network. By first calculating the cell-by-gene accessibility scores across annotated genes, we were able to generate lagged input matrices for the scATAC-seq dataset (Fig. 4a). Because the underlying gene-expression and gene-accessibility distributions are both zero-inflated^8^, the resulting coaccessibility matrices are qualitatively similar to gene-coexpression matrices.

**Figure 4:**
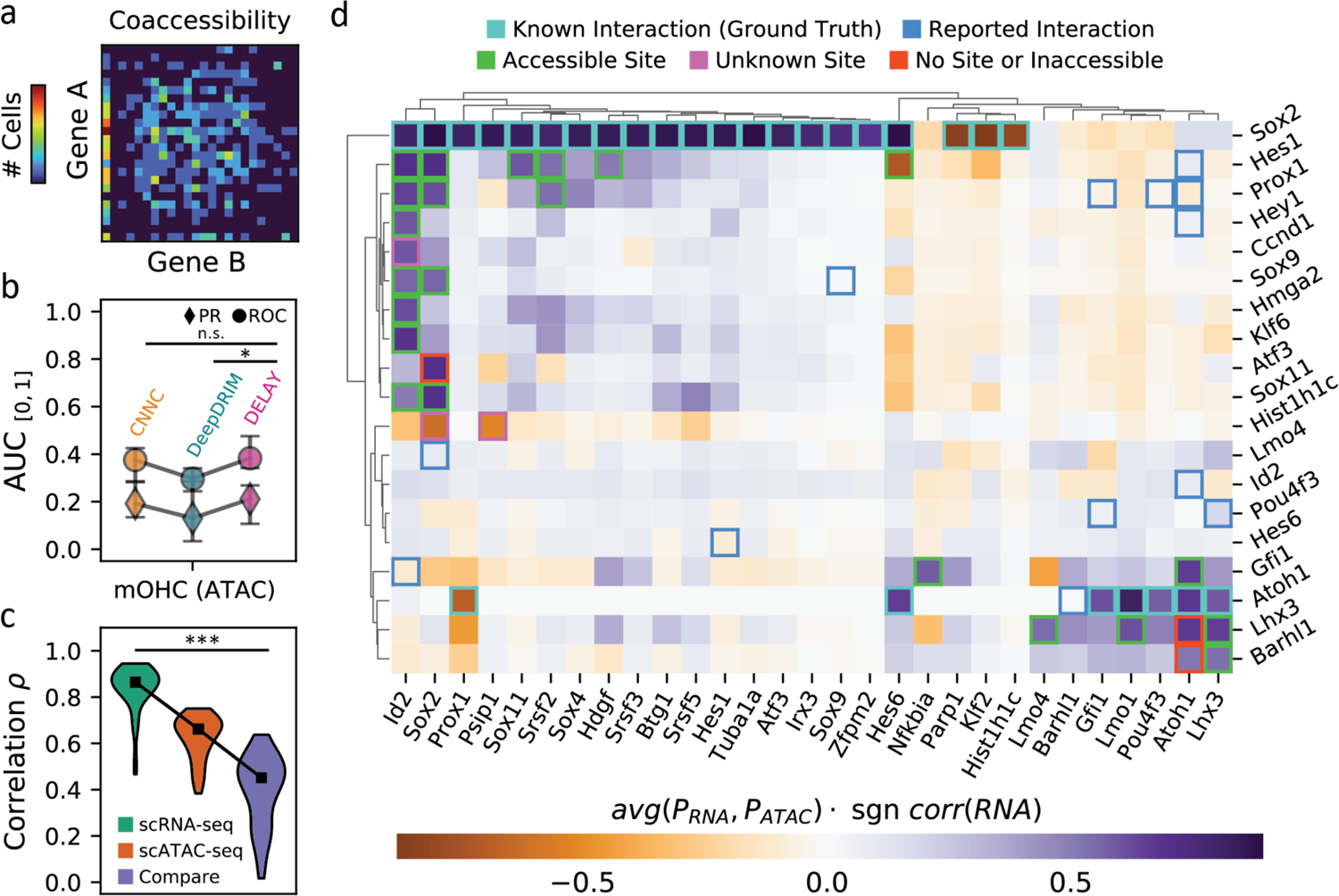
DELAY accurately predicts gene-regulatory interactions in the auditory hair cell network through multi-omic transfer learning. **a.** An example of an empirical joint-probability matrix from scATAC-seq trajectories. **b.** After fine-tuning the network on lagged scATAC-seq input matrices, DELAY performs comparably to other neural networks and to previous testing on hHep scRNA-seq data. **c.** Training DELAY separately on scRNA-seq and scATAC-seq datasets of hair cell development reveals a stronger correlation between predictions from cross-validated models (green and orange) than between datasets (purple) across the transcription factor-only network. **d.** Average gene-regulation probabilities *P* across the hair cell gene-regulatory network accurately predict interactions between transcription factors (rows) and targets (columns), when comparing known binding sites to open-chromatin peaks in the target genes’ enhancer sequences. Hierarchical clustering with WPGMA reveals distinct gene modules for prosensory genes (*Sox2*, *Id2*, *Hes1*, *Prox1*) and hair cell genes (*Atoh1*, *Pou4f3*, *Gfi1*, *Lhx3*, *Barhl1*). Up- or down-regulation was deduced from the correlation in the gene-expression data. Error bars and markers in **b** show the full range and average performance for each neural network across five cross-validated models. Markers in **c** show the median target gene-rank correlations across comparisons. The statistical significance in **b** was assessed with a one-sided Wilcoxon signed-rank test (*, *P*≤0.05) and in **c** with a Kruskal-Wallis test (***, *P*≤0.001).

To determine whether pseudotime-lagged gene-coaccessibility matrices contain features that indicate direct gene-regulatory relationships, we fine-tuned DELAY on the scATAC-seq matrices using available ground-truth examples collected from two cell type-specific datasets of Sox2 and Atoh1 target genes^32, 33^. Seventy percent of the ground-truth examples were used for fine-tuning, and the remaining 30 % were held out for validation (Fig. 4b). DELAY again performed slightly worse by the metric of area under the PR curve than by area under the ROC curve, indicating that the network is better at discriminating false positives than selecting true positives. Although DELAY did not outperform all other deep learning-based methods, the network’s performance was comparable to previous training on small fractions of the hHep scRNA-seq dataset, which suggests that gene-coaccessibility matrices are also useful for inferring direct gene-regulatory interactions.

Given that DELAY is well-optimized and benefits from additional training examples, we separately fine-tuned the network on all available Sox2 and Atoh1 ground-truth examples from an additional mOHC gene-expression dataset^30^. With previously determined hyperparameters for scRNA-seq data, training DELAY on two GPUs required 230 ± 1 min (mean ± SD) per cross-validated model. We then compared the target gene-rank correlations between the inferred transcription factor-only gene-regulatory networks to determine whether networks inferred from scRNA-seq data and scATAC-seq data were similar (Fig. 4c). Although we observed stronger correlations between predictions from dataset-specific, cross-validated models than between average predictions across datasets, the target gene-rank correlations between the two datasets were highly variable and the predictions of some transcription factors agreed better than others. Reasoning that the inferences with better agreement across datasets constituted the best predictions and highest-confidence interactions, we derived the consensus hair cell gene-regulatory network from the average predictions across both datasets. The resulting network consisted of 347 predicted transcription factor-target gene interactions with regulation probabilities greater than 0.5.

### DELAY identifies important genes, interactions, and modules in the hair cell gene-regulatory network

We used hierarchical clustering to group transcription factors and target genes in the transcription factor-only gene-regulatory network for hair cells by similarities in their predicted targets and regulators, respectively (Fig. 4d). This procedure revealed two distinct developmental modules corresponding to prosensory genes such as *Sox2*, *Id2*, *Hes1*, and *Prox1* and hair cell genes such as *Atoh1*, *Pou4f3*, *Gfi1*, *Lhx3*, and *Barhl1*. We sought to validate the predicted interactions by comparing the locations of known transcription factor-binding sites^34, 35^ to the locations of open-chromatin peaks^31^ within 50 kb of target genes’ transcription start sites. Twenty-two of 28 predicted interactions were confirmed by the accessibility of transcription-factor binding sites. Of the six remaining interactions, three were instances of predicted target-gene regulation by transcriptional cofactors lacking true DNA-binding domains. We additionally identified 13 reported interactions that were not detected by DELAY^31, 36–54^.

The most notable feature of the inferred gene-regulatory network for hair cells is that *Sox2* and *Id2*—in addition to the well-known roles of their products in the maintenance of a prosensory-cell fate and the regulation of target genes—are themselves the targets of a wide variety of transcription factors including Sox proteins, homeobox factors, zinc-finger proteins, and Notch effectors. Eleven of the 22 interactions confirmed by our binding-site accessibility analysis represented direct regulation of *Sox2* and *Id2*, including mutually activating interactions between *Sox2* and *Sox9*, *Sox11*, *Prox1*, and *Hes1*, and mutual inhibition with *Hist1h1c*. One other study has suggested that Hes1 directly regulates *Sox2*^55^.

Another notable feature of the inferred hair cell network is that the LIM-homeobox transcription factor Lhx3 regulates its own expression as well as that of two other LIM-only transcription factors (Lmo4 and Lmo1). This result implies a role for Lhx3 in maintaining the later expression of these important early *Sox2* inhibitors^31, 47, 49^. Other key features of the network include *Sox11* regulation by Hes1, *Nfkbia* regulation by Gfi1, and regulation of the splicing-factor gene *Srsf2* by the products of three different prosensory genes. Srsf2 was recently predicted to play a role in the splicing of an important deafness gene in humans^56^.

### Discriminative motif analysis of target-gene enhancer sequences enables *de novo* discovery of DNA-binding motifs

DELAY permits a complementary approach to typical gene-regulatory inference workflows such as SCENIC^57^ that use *cis*-regulatory sequences to identify and discard false-positive interactions resulting from indirect gene regulation^11, 30, 31^. Because these methods rely on databases of known DNA-binding motifs, they necessarily overlook predictions for cofactors and putative transcription factors with unknown binding sequences. Instead of using *cis*-regulatory sequences to refine our initial predictions, we introduced enhancer-sequence information *post hoc* to perform discriminative motif analysis within the enhancers of predicted target genes in the hair cell network (Fig. 5a, Supplementary Fig. 6). Notably, we identified enriched motifs that closely resembled known DNA-binding motifs for nine of eleven transcription factors with at least one predicted target gene in the transcription factor-only hair cell network (Supplementary Table 2).

**Figure 5:**
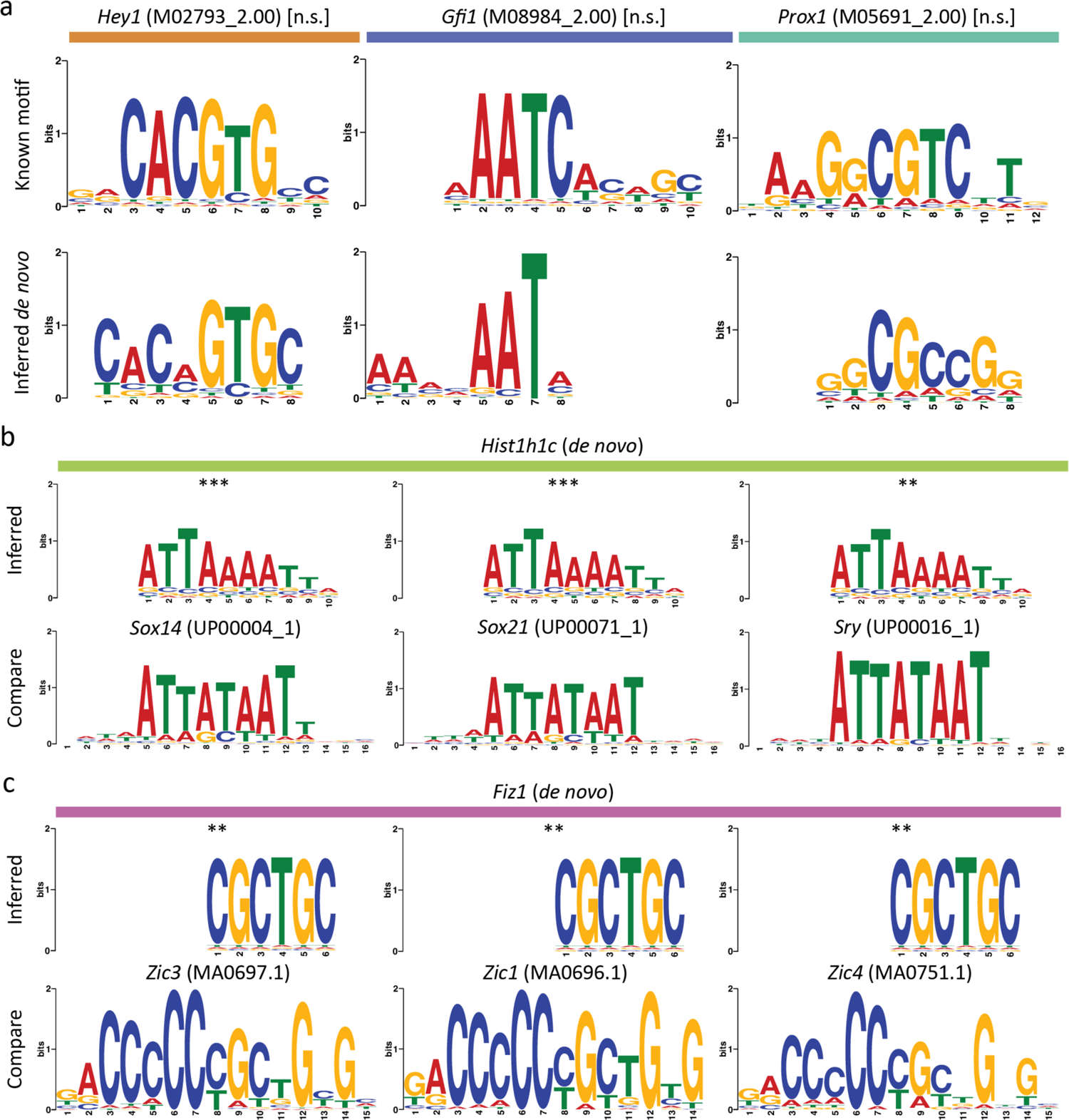
Accurate target-gene predictions enable *de novo* discovery of DNA-binding motifs. **a.** Three examples of motifs enriched in the enhancer sequences of predicted target genes (bottom row) which closely resemble known DNA-binding motifs (top row) for transcription factors with at least one predicted target gene in the transcription factor-only hair cell gene-regulatory network. **b, c.** Sequences for cofactors or putative transcription factors may represent novel DNA-binding motifs or indicate sequence-specific interactions through multi-protein complexes. Motifs for two hair cell genes closely match known DNA-binding sequences, suggesting that the linker histone Hist1h1c (**b**, top row) forms complexes with SoxB2/A factors (**b**, bottom row), and that the putative C2H2 zinc-finger protein Fiz1 (**c**, top row) recognizes motifs similar to Zic-family C2H2 zinc-finger proteins (**c**, bottom row). The statistical significance of each motif alignment was estimated using the cumulative density function of all possible comparisons of known motifs across enriched sequences for **a,** or inferred motifs across all database motifs for **b** and **c** (**, *P*≤0.01; ***, *P*≤0.001).

We additionally sought to predict the mechanisms by which several identified hair cell cofactors accomplish sequence-specific gene regulation in the absence of true DNA-binding domains. Specifically, we compared sequences enriched in the enhancers of cofactors’ predicted targets to several databases of known DNA-binding motifs to identify the cofactors’ DNA-binding partners (so-called “guilt by association”). Through this analysis, we were able to determine that the most significantly enriched sequence in the enhancers of the histone Hist1h1c’s predicted target genes closely matched known binding motifs for several SoxB2/A transcription factors (Fig. 5b). In addition, a sequence enriched in the enhancers of cyclin Ccnd1’s targets closely matched known motifs for the Pbx family of homeobox transcription factors (Supplementary Fig. 7). These cofactors might form complexes with the identified transcription factors to regulate their target genes in hair cells.

As a final demonstration of DELAY’s high-confidence predictions, we used the fine-tuned scRNA-seq models to generate regulatory predictions for transcription factors expressed in at least 20 % of developing hair cells (Supplementary Table 3). After considering these additional transcription factors, we identified the putative C2H2 zinc-finger protein Fiz1 as a likely regulator of *Sox2* and *Hes6*. Discriminative motif analysis of these genes’ enhancer sequences (Fig. 5c) uncovered a likely Fiz1 consensus DNA-binding sequence (5’-CGCTGC-3’) similar to that of other *Sox2* regulators from the Zic family of C2H2 zinc-finger proteins^58^.

## Discussion

Building upon several deep learning-based methods^18, 22^, we have demonstrated that combining fully-supervised deep learning with joint-probability matrices of pseudotime-lagged single-cell trajectories can overcome certain limitations of current Granger causality-based methods of gene-regulatory inference^9, 12^, such as their inability to infer cyclic regulatory motifs. Moreover, although our convolutional neural network DELAY remains sensitive to the ordering of single cells in pseudotime, the network can accurately infer direct gene-regulatory interactions from both timecourse and snapshot datasets, unlike many supervised methods that rely strongly on the number of available time-course samples^15, 16^. We suspect that DELAY is sensitive to the specific ordering of adjacent cells in trajectories because pseudotime inference methods such as Slingshot^6^ infer lineages from minimum spanning trees that directly depend on cell-to-cell similarities in gene-expression values^7^.

We have presented a multi-omic paradigm for fine-tuning DELAY on both gene-expression and chromatin-accessibility datasets for the development of auditory hair cells. The network’s accurate predictions allowed *de novo* inferences of known and unknown DNA-binding motifs, establishing DELAY as a complementary approach to common methods of gene-regulatory inference. Our method’s high-confidence predictions across the hair cell gene-regulatory network also support it as an attractive option for experimentalists with limited resources to predict true gene-regulatory relationships from complex, large-scale gene-regulatory networks while avoiding spurious, indirect interactions often introduced through unsupervised methods^14^.

We have additionally identified with high confidence interactions between cofactors and target genes in the regulatory network of hair cells that methods such as SCENIC^57^ would have overlooked. Using “guilt by association,” we predicted that the cyclin Ccnd1—a known transcriptional cofactor found at the enhancers of several *Id* genes during retinal development^59^— forms complexes with the Pbx family of homeobox transcription factors, of which at least one is a known target of Prox1 in the inner ear^60^. Moreover, we predicted that the retina-specific linker histone Hist1h1c^61^ forms complexes with the SoxB2 family of transcription factors, which have known antagonistic effects on *Sox2* expression in hair cells^62^. Finally, we predicted that the putative hair cell-specific C2H2 zinc-finger protein Fiz1—which is also expressed during retinal development^63^—is a regulator of *Sox2* in hair cells and identified its likely DNA-binding sequence in the enhancers of predicted target genes.

Believing that DELAY will be a valuable resource to the community, we have provided an open-source implementation of the algorithm as well as pre-trained model weights for subsequent fine-tuning on new single-cell datasets and partial ground-truth labels. Although we chose to fine-tune DELAY on cell type-specific ChIP-seq data, future studies may choose to fine-tune DELAY on curated interactions from databases or from gain-or loss-of-function experiments, especially in the absence of ChIP-seq data^14^. These additional interactions can also supplement smaller ground-truth datasets, such as those of Sox2 and Atoh1 target genes, to mitigate false-negative predictions. We believe the modest computational cost of training new models of DELAY will prove a worthwhile investment for future investigations into complex gene-regulatory networks.

## Methods

### Preparing single-cell RNA-seq datasets to train and test the convolutional neural network

To train our convolutional neural network, we used scRNA-seq datasets from two timecourse studies of endodermal specification of human embryonic stem cells^19^ and mouse embryonic stem cells^20^ and an *in vivo* snapshot study of erythroid (E), granulocyte-monocyte (GM), and lymphoid (L) specification in mouse hematopoietic stem cells^21^. We additionally employed a dataset from a timecourse study of the differentiation of human hepatocytes from induced pluripotent stem cells to test our network^28^. For each of these studies, we collected the normalized gene-expression values, pseudotime values, and ground-truth networks from BEELINE’s supplementary data files^14, 64^. In BEELINE, Pratapa *et al*. collected the normalized gene-expression values from the original studies or obtained the values themselves by log-transforming the transcripts per kilobase million. Moreover, those authors used Slingshot^6^ to infer the pseudotime values separately for each dataset, orienting the inferred trajectories by known experimental timepoints or precursor-cell types and lineages. Finally, Pratapa *et al*. selected genes with differential expression across pseudotime by applying generalized additive models (GAMs) to compute gene variances and their associated *P* values. To train DELAY, we utilized their gene variances and corrected *P* values to select differentially expressed transcription factors (*P*_adjusted_<0.01) and 500 additional genes with the highest variance for each dataset. We also collected the ground-truth labels from BEELINE’s supplementary data files, which contained tables of transcription factor-target gene interactions curated from cell type-specific ChIP-seq experiments in the ENCODE^65^, ChIP-Atlas^66^, and ESCAPE^67^ databases. Table 1 provides details for each dataset including descriptions of the ground-truth networks.

We also characterized our network on modified datasets with shuffled pseudotime, fewer cells, and additional gene-dropout noise. We shuffled pseudotime values across single-cell trajectories using the np.random.normal function in NumPy (v1.20.2) to select the indices of single cells either leading or lagging successive cells in each trajectory at some length scale *σ* before swapping the cells’ positions in pseudotime. We used np.random.choice in NumPy to select the indices of single cells to retain when down-sampling the number of cells in a dataset. Lastly, we induced additional gene-dropout noise in datasets by setting the gene-expression values to zero with a probability of *p* for the bottom *p* percent of cells expressing each gene in a given dataset.

### Generating examples of transcription factor-target gene pairs from single-cell datasets

For each dataset, we generated mini-batches of 512 transcription factor-target gene-pair examples by first enumerating all possible combinations of differentially expressed transcription factors and potential target genes—including both transcription factors and highly varying genes. We then generated aligned and pseudotime-lagged gene-coexpression matrices for up to five lags with the following configurations as separate input channels: transcription factor-target, transcription factor-transcription factor, target-target, transcription factor-neighbor (for two neighbors), and target-neighbor (for two neighbors). We concatenated the input channels for each example to form three-dimensional stacks of input matrices with dimensions 42×32×32 (channels x height x width). For each gene, we used the np.correlate function in NumPy to select the transcription factors with the highest absolute cross-correlation at a maximum offset of five pseudotime lags to use as neighbor genes. After preparing mini-batches for each dataset, we used different integer values to seed the random_split function in PyTorch (v1.8.1) to produce pseudorandom, reproducible training (70 %) and validation (30 %) splits across each dataset for cross validation. We later used the same validation splits to characterize our network on augmented coexpression matrices and matrices generated from modified single-cell datasets. To augment gene-coexpression matrices, we masked either specific regions across all input channels, or entire input channels corresponding to specific pseudotime lags, by setting all of the relevant pixel values to zero. We replaced features after motif-specific neighbor-gene ablations by selecting new highly correlated genes to replace neighbors involved in the higher-order gene-regulatory interactions.

### Constructing a convolutional neural network to classify lagged gene-coexpression matrices

We designed a convolutional neural network based on an inverted VGGnet^68^ that uses five convolutional layers to first expand the input to 1024 channels, then successively halve the number of channels to 64 before classifying examples as either true regulation or no interaction with a fully-connected linear layer. Each convolutional layer sums over the two-dimensional cross-correlations (★) of 3×3 kernels and input channels *i* to find the features for a given output channel *j*, as shown in Equation 1:

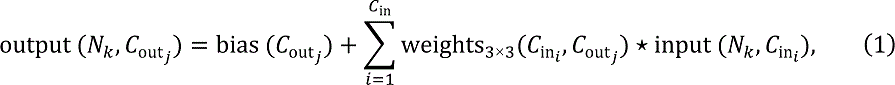

in which the weights and bias are trainable parameters, *N* is the mini-batch size, *C* is the number of channels, *C*_out_ = *C*_in_⁄2 for layers two through five, and all convolutions are zero-padded at the edges of matrices. To preserve the gradient flow during training, we used leaky rectified linear units (ReLUs) with a negative slope of 0.2 (Equation 2) as our nonlinear activation function after each convolutional layer:

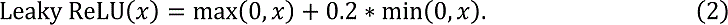

As shown in Equation 3, we additionally used 2×2, unpadded max-pooling layers between convolutional layers to identify important features and down-sample activation maps:

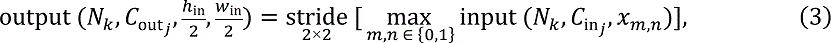

in which *x* represents the 2×2 input windows and ℎ, *W* are the height and width of the input channel *j*. After the final convolutional layer, we used global-average pooling (Equation 4) to reduce the remaining 64 feature maps *j* to a single, 64-dimensional latent vector x:

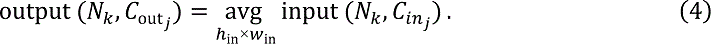

We lastly used a fully-connected, linear layer (Equation 5) with a sigmoid activation function (Equation 6) to generate gene-regulation probabilities:

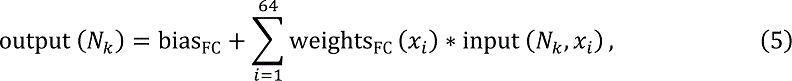

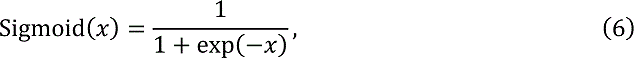

where a probability of *P* > 0.5 indicates a true gene-regulatory interaction for the given gene-pair example.

### Training and fine-tuning the network on pseudotime-lagged gene-coexpression matrices

We used PyTorch’s implementation of stochastic gradient descent (SGD) to optimize our network with respect to the class-weighted binary cross-entropy loss, summed across each mini-batch and scaled by the overall batch size 512, as shown in Equation 7:

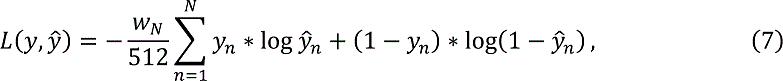

in which *y* and ŷ are the target and predicted values (respectively), *W*_E_ is the fraction of true examples in the mini-batch, and *N* is the size of the current mini-batch (which might be less than 512). Prior to optimization, we used He initialization with uniform priors to set the weights for all convolutional and linear layers. We used a learning rate (LR) of 0.5 to train each model for up to 100 epochs, validating performance after each epoch, and stopping training after ten or more epochs without an improvement in the average validation metric of areas under the curves.

For fine-tuning, we increased the LR and number of epochs over several orders of magnitude when training models on smaller fractions of the testing dataset. To a lesser extent, we also adjusted the LR and number of epochs when training models on different configurations of lagged matrices, neighbor-gene matrices, and matrices of different sizes to improve training speed and stability. When fine-tuning the network on scATAC-seq matrices, we used mini-batches with 24 examples each and trained for 50 to 100 epochs. We used two Nvidia RTX 8000 GPUs to train each model.

### Comparing DELAY to other top-performing gene-regulatory inference methods

We compared the performance of our neural network to two other convolutional neural networks, as well as the top six best-performing methods identified in a previous benchmarking study. Using BEELINE, we inferred the gene-regulatory networks for all training and testing datasets using the tree-based methods GENIE3^23^ and GRNBoost2^24^, the mutual information-based method PIDC^25^, and the partial-correlation and regression-based methods PPCOR^26^, SCODE^27^, and SINCERITIES^8^. We utilized the best parameter values identified in BEELINE for the partial-correlation and regression-based methods. For the two deep learning-based methods (CNNC^22^ and DeepDRIM^18^), we used the original studies to reconstruct the neural networks in PyTorch before training five cross-validated models each. In addition to training both networks on pseudotime-aligned gene-coexpression matrices for primary transcription factor-target gene pairs, we also trained DeepDRIM on ten neighbor-gene matrices per gene. Neither network was trained on pseudotime-lagged gene-coexpression matrices.

### Analyzing enrichment of gene-ontology terms across correctly inferred gene-pair examples

We used Enrichr^69^ (https://maayanlab.cloud/Enrichr/) to analyze GO-term enrichment across correctly inferred transcription factors for terms related to InterPro DNA-binding domains^70^. We used a Fisher’s exact test with Benjamini-Hochberg correction for multiple-hypothesis testing (*P*_adjusted_<0.05) to assess the statistical significance for each GO term.

### Preparing single-cell multi-omics datasets to infer the hair cell gene-regulatory network

We used two single-cell datasets from developing mouse outer hair cells^30, 31^ to predict the consensus hair cell gene-regulatory network. First, we collected the normalized gene-expression values from the original scRNA-seq study^30^, then used reciprocal PCA to integrate single cells across four timepoints of sensory-epithelium development into a single dataset in Seurat^71^ (v3.1.4). Then, we used Slingshot^6^ (v1.0.0) to infer pseudotime values across the outer hair cell trajectory and applied LOESS regressions and GAMs to select the genes that were differentially expressed across both the sensory epithelium and pseudotime, respectively. We again selected all differentially expressed transcription factors (*P*<0.01 after Bonferroni correction for multiple-hypothesis testing) and 500 additional genes with the highest variance, and we used the integrated gene-expression assay in Seurat to generate the corresponding gene-coexpression matrices. In a separate analysis, we broadened our selection criteria to encompass all genes that were differentially expressed in at least 20 % of the outer hair cells, regardless of their expression across the full sensory epithelium.

For the scATAC-seq dataset, we first collected the cell-by-peak chromatin-accessibility data from the original study^31^. Then, we used the function createGmatFromMat in SnapATAC^72^ (v1.0.0) to calculate the cell-by-gene accessibility scores as the counts of each 5 kb bin per gene in the UCSC mouse genome mm10^73^ (TxDb.Mmusculus.UCSC.mm10.knownGene; v3.4.4) that contained at least one open-chromatin peak for a given cell. These values were then log-normalized. We again used Slingshot to infer pseudotime values across the outer hair cell trajectory and applied GAMs to the raw accessibility counts to select the differentially accessible genes across pseudotime. For the inferred scATAC-seq network, we selected all differentially accessible transcription factors and variable genes (*P*_adjusted_<0.01) that were also differentially expressed along the scRNA-seq trajectory. Because the scATAC-seq dataset had fewer single cells than the scRNA-seq dataset, we used 24 fixed-width bins in each dimension to generate the corresponding gene-coaccessibility matrices prior to fine-tuning.

### Discriminative motif analysis to discover *de novo* transcription factor binding motifs

We used the UCSC mouse genome mm10 and twoBitToFa to download the 100 kb enhancer sequences spanning 50 kb upstream and downstream of all target genes’ transcription start sites in the transcription factor-only hair cell gene-regulatory network, then divided them into 100 bp fragments and sorted the resulting sequences into primary (predicted targets) and control (predicted no interaction) groups for each transcription factor with at least one target gene in the network. We then used STREME^74^ (v5.4.1) to perform discriminative motif analysis with a *P* value threshold of 0.05 to identify enriched motifs in predicted target genes’ enhancers, which we then compared to either the transcription factors’ known binding motifs from CIS-BP^34^, or to all motifs from the JASPAR^75^, UniPROBE^76^ (mouse), and Jolma et al. (2013)^77^ databases using TOMTOM^78^ (v5.4.1) with an *E* value threshold of 10 for significant alignments.

## Data availability

The processed experimental files for all single-cell datasets used in this study are available on Zenodo at https://doi.org/10.5281/zenodo.5711739; Table 1 lists the Gene Expression Omnibus (GEO) accession numbers for each dataset. The saved model weights for DELAY are available on Zenodo at https://doi.org/10.5281/zenodo.5711792. All experimental logs from this study are available at https://tensorboard.dev/experiment/RBVBetLMRDiEvO7sBl452A.

## Code availability

We have provided an open-source implementation of DELAY in PyTorch with listed requirements and documentation at https://github.com/calebclayreagor/DELAY.

## Acknowledgements

The authors would like to acknowledge Junyue Cao, Viviana Risca, Christina Leslie, Adrian Jacobo, Agnik Dasgupta, Emily Atlas, and other members of the Laboratory of Sensory Neuroscience for helpful discussions and comments on the manuscript. C.C.R. is supported by National Science Foundation Graduate Research Fellowship Grant No. 1946429. A.J.H is an Investigator of Howard Hughes Medical Institute.

## Author contributions

C.C.R. and N.V. conceived the study, and all authors designed the analysis. C.C.R. carried out all experiments and analysis and wrote the paper. A.J.H. supervised the project, and all the authors edited the paper.

## Competing interests

The authors declare no competing interests.

---

**Supplementary Figure 1:**
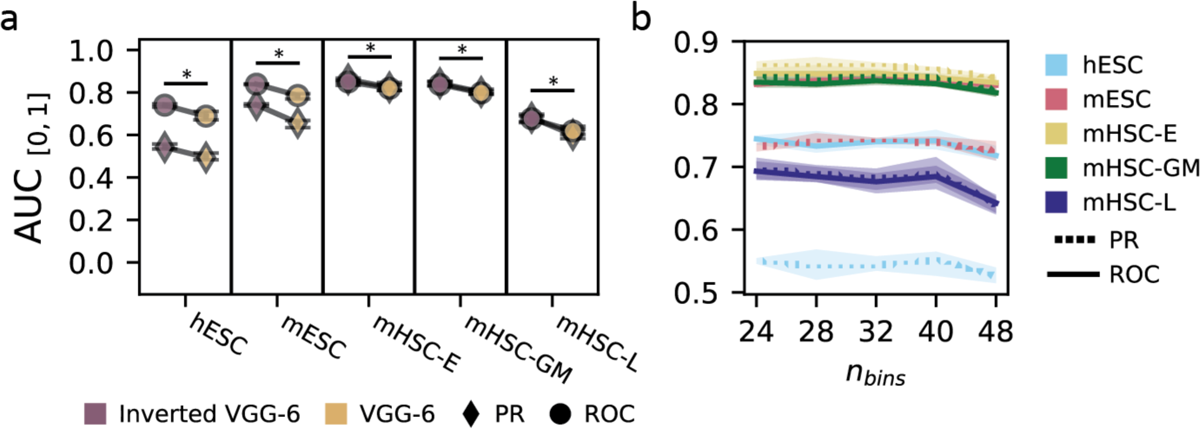
The inverted VGG-6 network outperforms a conventional VGG-6. **a.** Utilizing a more expressive feature space at the top of the network allows the inverted VGG-6 to outperform a conventional VGG-6 network when trained on the same input data. **b.** Performance of the inverted VGG-6 network is robust across input-matrix sizes. Error bars and markers in **a** and **b** show the full range and average performance across five cross-validated models. The statistical significance in **a** was assessed using a one-sided Wilcoxon signed-rank test (*, *P*≤0.05).

**Supplementary Figure 2:**
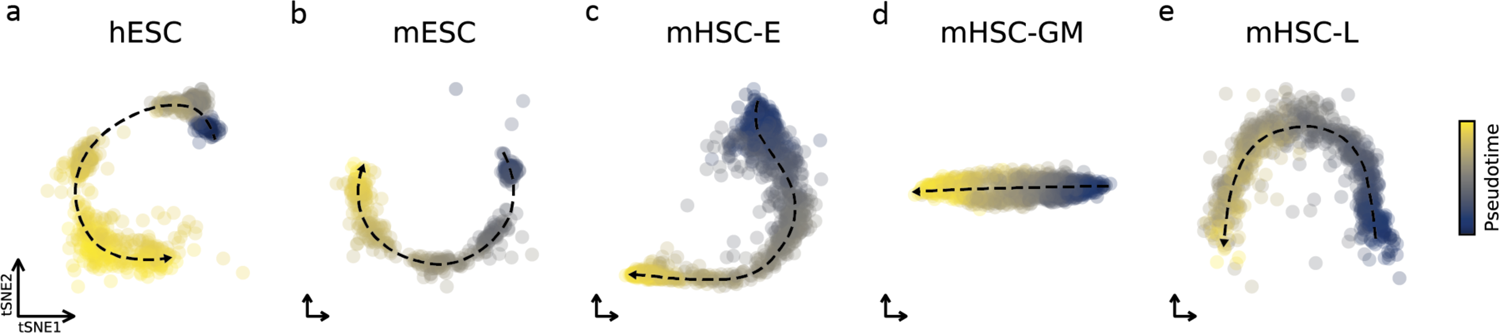
Inferred pseudotime trajectories for the five training datasets. Two-dimensional embeddings of the trajectories for human embryonic stem cells (**a**), mouse embryonic stem cells (**b**), and mouse hematopoietic stem cells (**c-e**) were generated from the differentially expressed genes identified in BEELINE and colored by the inferred Slingshot pseudotime values.

**Supplementary Figure 3:**
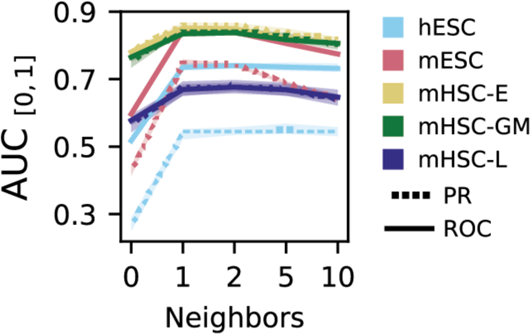
Neighbor-gene matrices improve the neural network’s performance. Training the network on joint-probability matrices of up to two highly correlated neighbor genes increases performance across all datasets. Error bars and lines show the full range and average performance across five cross-validated models.

**Supplementary Figure 4:**
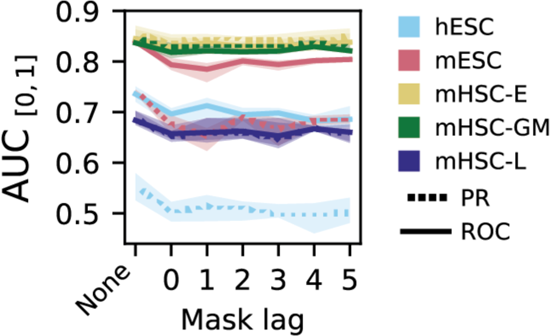
Augmenting input matrices decreases the network’s performance. Even though performance increases the most when training DELAY on input matrices from up to a single pseudotime lag, testing on channel-masked inputs suggests that the network relies on all available inputs, even gene-coexpression matrices from pseudotime lags two through five. Error bars and lines show the full range and average performance across five cross-validated models.

**Supplementary Figure 5:**
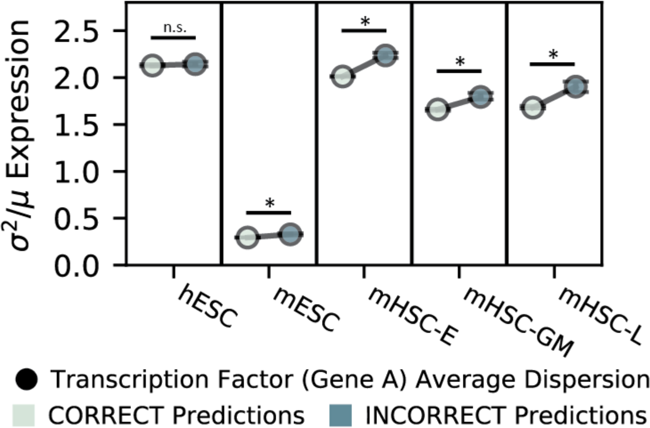
DELAY performs best on transcription factors with stable gene expression. An analysis of statistical dispersion across transcription factors shows that on average correctly inferred examples have lower variance-to-mean ratios than those of incorrect predictions. Error bars and markers show the full range and mean values across predictions from five cross-validated models. The statistical significance was assessed using a one-sided Wilcoxon signed-rank test (*, *P*≤0.05).

**Supplementary Figure 6:**
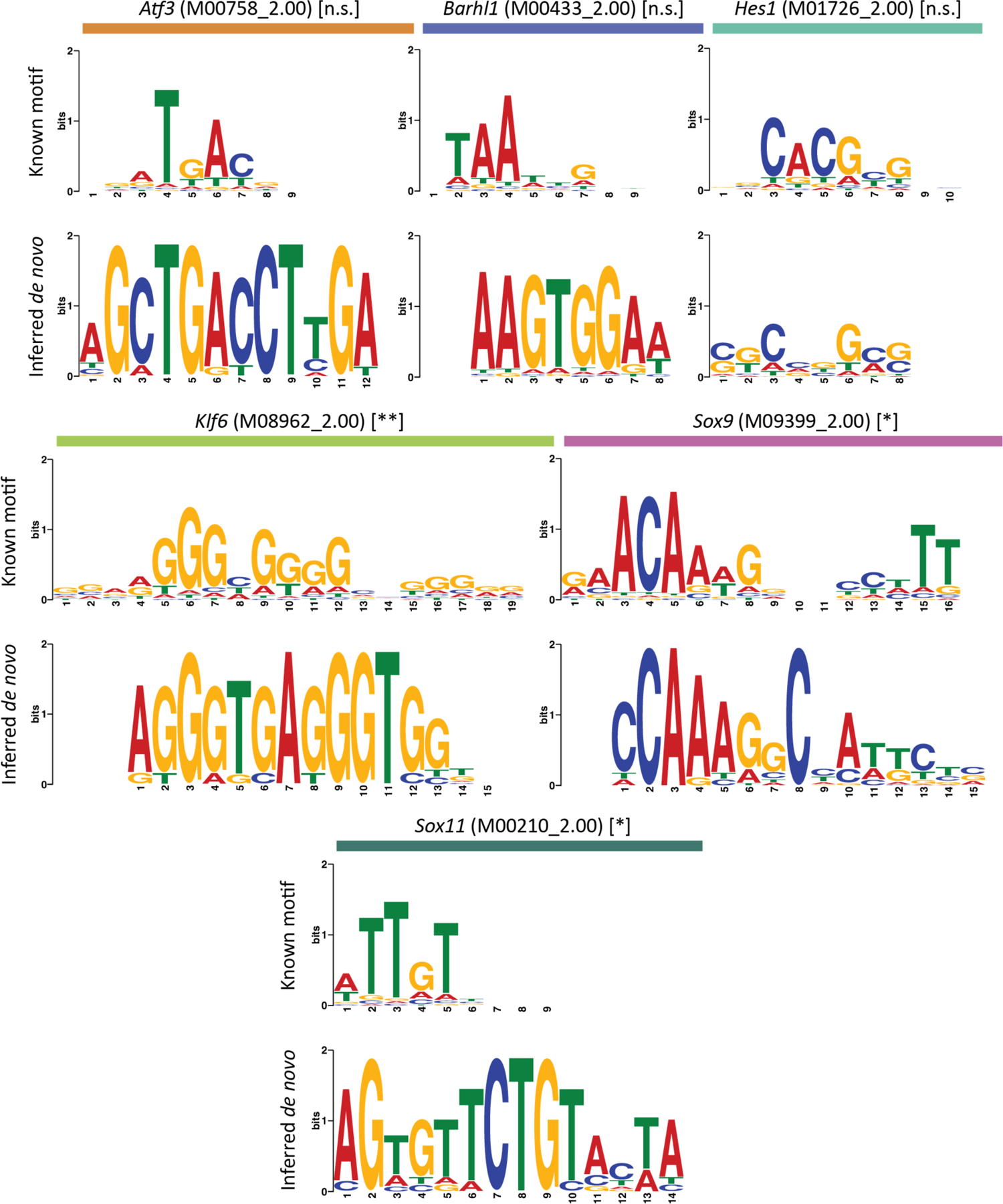
Discriminative motif analysis corroborates gene-regulatory predictions. Six additional examples of enriched motifs in the enhancer sequences of predicted target genes which closely resemble known DNA-binding motifs for transcription factors with at least one predicted target gene in the transcription factor-only hair cell gene-regulatory network. The statistical significance of each motif alignment was estimated using the cumulative density function of all possible comparisons of known motifs across enriched sequences (*, *P*≤0.05; **, *P*≤0.01).

**Supplementary Figure 7:**
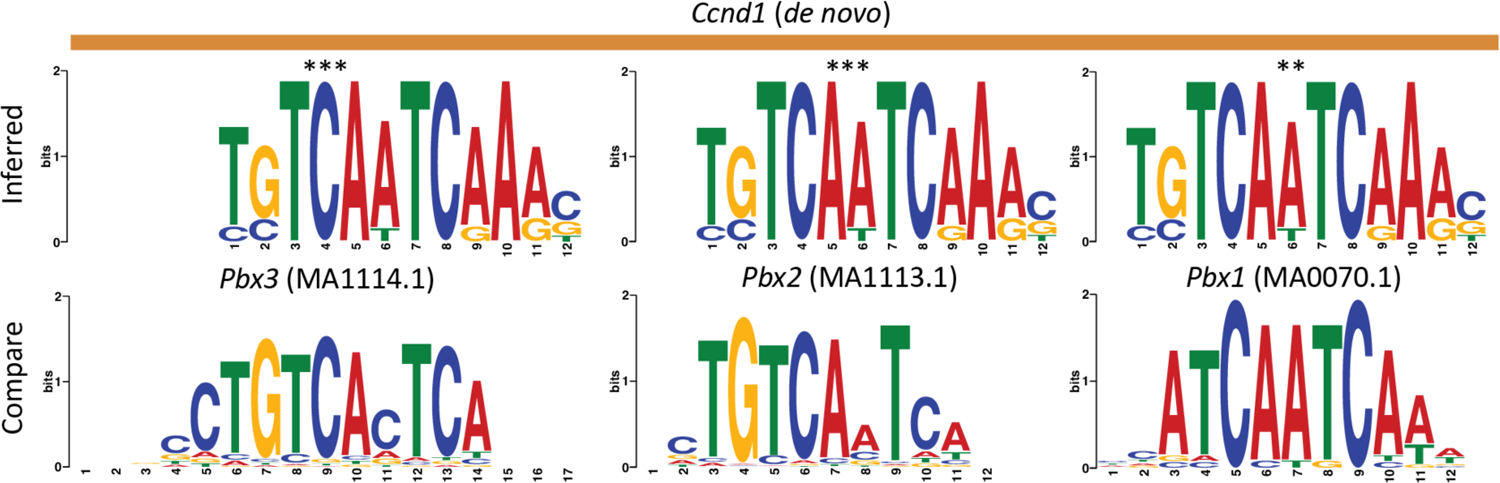
An enriched motif in the enhancers of Ccnd1’s predicted target genes suggests a mechanism for cofactor-target interactions. An enriched motif in the enhancer sequences of cyclin Ccnd1’s predicted target genes (top row) suggests that the putative cofactor forms complexes with Pbx-family homeobox transcription factors (bottom row) to regulate its target genes in a sequence-specific manner. The statistical significance of each motif alignment was estimated using the cumulative density function of all possible comparisons of the inferred motif across all database motifs (**, *P*≤0.01; ***, *P*≤0.001).

## Notes

https://doi.org/10.5281/zenodo.5711739

https://doi.org/10.5281/zenodo.5711792

https://tensorboard.dev/experiment/RBVBetLMRDiEvO7sBl452A

https://github.com/calebclayreagor/DELAY

### Competing Interest Statement

The authors have declared no competing interest.

### Summary of Updates

Corrected Gene Expression Omnibus Accession Numbers in Table 1

## References

1. Ocone, A., Haghverdi, L., Mueller, N. S. & Theis, F. J. Reconstructing gene regulatory dynamics from high-dimensional single-cell snapshot data. Bioinformatics 31, i89–i96 (2015).

2. Qian, J., Dolled-Filhart, M., Lin, J., Yu, H. & Gerstein, M. Beyond synexpression relationships: local clustering of time-shifted and inverted gene expression profiles identifies new, biologically relevant interactions. J. Mol. Biol. 314, 1053–1066 (2001).

3. Bar-Joseph, Z., Gitter, A. & Simon, I. Studying and modelling dynamic biological processes using time-series gene expression data. Nat. Rev. Genet. 13, 552–564 (2012).

4. Ding, J., Sharon, N. & Bar-Joseph, Z. Temporal modelling using single-cell transcriptomics. Nat. Rev. Genet. (2022) doi:10.1038/s41576-021-00444-7.

5. Tritschler, S. et al. Concepts and limitations for learning developmental trajectories from single cell genomics. Development 146, dev170506 (2019).

6. Street, K. et al. Slingshot: Cell lineage and pseudotime inference for single-cell transcriptomics. BMC Genomics 19, 477 (2018).

7. Trapnell, C. et al. The dynamics and regulators of cell fate decisions are revealed by pseudotemporal ordering of single cells. Nat. Biotechnol. 32, 381–386 (2014).

8. Papili Gao, N., Ud-Dean, S. M. M., Gandrillon, O. & Gunawan, R. SINCERITIES: Inferring gene regulatory networks from time-stamped single cell transcriptional expression profiles. Bioinformatics 34, 258–266 (2018).

9. Deshpande, A., Chu, L.-F., Stewart, R. & Gitter, A. Network inference with Granger causality ensembles on single-cell transcriptomics. Cell Rep. 38, 110333 (2022).

10. Finkle, J. D., Wu, J. J. & Bagheri, N. Windowed Granger causal inference strategy improves discovery of gene regulatory networks. Proc. Natl. Acad. Sci. 115, 2252–2257 (2018).

11. Michailidis, G. & d’Alché-Buc, F. Autoregressive models for gene regulatory network inference: Sparsity, stability and causality issues. Math. Biosci. 246, 326–334 (2013).

12. Zhang, Y., Chang, X. & Liu, X. Inference of gene regulatory networks using pseudo-time series data. Bioinformatics btab099 (2021) doi:10.1093/bioinformatics/btab099.

13. Qiu, X. et al. Inferring causal gene regulatory networks from coupled single-cell expression dynamics using scribe. Cell Syst. 10, 265–274.e11 (2020).

14. Pratapa, A., Jalihal, A. P., Law, J. N., Bharadwaj, A. & Murali, T. M. Benchmarking algorithms for gene regulatory network inference from single-cell transcriptomic data. Nat. Methods 17, 147–154 (2020).

15. Yuan, Y. & Bar-Joseph, Z. Deep learning of gene relationships from single cell time-course expression data. Brief. Bioinform. 22, bbab142 (2021).

16. Xu, Y., Chen, J., Lyu, A., Cheung, W. K. & Zhang, L. dynDeepDRIM: a dynamic deep learning model to infer direct regulatory interactions using single cell time-course gene expression data. http://biorxiv.org/lookup/doi/10.1101/2021.08.28.458048 (2021) doi:10.1101/2021.08.28.458048.

17. Granger, C. W. J. Investigating causal relations by econometric models and cross-spectral methods. Econometrica 37, 424–38 (1969).

18. Chen, J. et al. DeepDRIM: A deep neural network to reconstruct cell-type-specific gene regulatory network using single-cell RNA-seq data. Brief. Bioinform. 22, (2021).

19. Chu, L.-F. et al. Single-cell RNA-seq reveals novel regulators of human embryonic stem cell differentiation to definitive endoderm. Genome Biol. 17, (2016).

20. Hayashi, T. et al. Single-cell full-length total RNA sequencing uncovers dynamics of recursive splicing and enhancer RNAs. Nat. Commun. 9, 619 (2018).

21. Nestorowa, S. et al. A single-cell resolution map of mouse hematopoietic stem and progenitor cell differentiation. Blood 128, e20–e31 (2016).

22. Yuan, Y. & Bar-Joseph, Z. Deep learning for inferring gene relationships from single-cell expression data. Proc. Natl. Acad. Sci. 116, 27151–27158 (2019).

23. Huynh-Thu, V. A., Irrthum, A., Wehenkel, L. & Geurts, P. Inferring regulatory networks from expression data using tree-based methods. PLoS ONE 5, e12776 (2010).

24. Moerman, T. et al. GRNBoost2 and Arboreto: Efficient and scalable inference of gene regulatory networks. Bioinformatics 35, 2159–2161 (2019).

25. Chan, T. E., Stumpf, M. P. H. & Babtie, A. C. Gene regulatory network inference from single-cell data using multivariate information measures. Cell Syst. 5, 251–267.e3 (2017).

26. Kim, S. ppcor: An R package for a fast calculation to semi-partial correlation coefficients. Commun. Stat. Appl. Methods 22, 665–674 (2015).

27. Matsumoto, H. et al. SCODE: An efficient regulatory network inference algorithm from single-cell RNA-Seq during differentiation. Bioinformatics 33, 2314–2321 (2017).

28. Camp, J. G. et al. Multilineage communication regulates human liver bud development from pluripotency. Nature 546, 533–538 (2017).

29. Alon, U. Network motifs: theory and experimental approaches. Nat. Rev. Genet. 8, 450– 461 (2007).

30. Kolla, L. et al. Characterization of the development of the mouse cochlear epithelium at the single cell level. Nat. Commun. 11, 2389 (2020).

31. Wang, S., Lee, M. P., Jones, S., Liu, J. & Waldhaus, J. Mapping the regulatory landscape of auditory hair cells from single-cell multi-omics data. Genome Res. gr.271080.120 (2021) doi:10.1101/gr.271080.120.

32. Kwan, K. Y., Shen, J. & Corey, D. P. C-MYC transcriptionally amplifies SOX2 target genes to regulate self-renewal in multipotent otic progenitor cells. Stem Cell Rep. 4, 47–60 (2015).

33. Cai, T. et al. Characterization of the transcriptome of nascent hair cells and identification of direct targets of the Atoh1 transcription factor. J. Neurosci. 35, 5870–5883 (2015).

34. Weirauch, M. T. et al. Determination and inference of eukaryotic transcription factor sequence specificity. Cell 158, 1431–1443 (2014).

35. Grant, C. E., Bailey, T. L. & Noble, W. S. FIMO: scanning for occurrences of a given motif. Bioinformatics 27, 1017–1018 (2011).

36. Zheng, J. L., Shou, J., Guillemot, F., Kageyama, R. & Gao, W. Q. Hes1 is a negative regulator of inner ear hair cell differentiation. Development 127, 4551–4560 (2000).

37. Zine, A. et al. Hes1 and Hes5 activities are required for the normal development of the hair cells in the mammalian inner ear. J. Neurosci. 21, 4712–4720 (2001).

38. Ikeda, R., Pak, K., Chavez, E. & Ryan, A. F. Transcription factors with conserved binding sites near ATOH1 on the POU4F3 gene enhance the induction of cochlear hair cells. Mol. Neurobiol. 51, 672–684 (2015).

39. Du, X. et al. Regeneration of mammalian cochlear and vestibular hair cells through Hes1/Hes5 modulation with siRNA. Hear. Res. 304, 91–110 (2013).

40. Kirjavainen, A. et al. Prox1 interacts with Atoh1 and Gfi1, and regulates cellular differentiation in the inner ear sensory epithelia. Dev. Biol. 322, 33–45 (2008).

41. Yu, K. S. et al. Development of the mouse and human cochlea at single cell resolution. 739680 (2019).

42. Benito-Gonzalez, A. & Doetzlhofer, A. Hey1 and Hey2 control the spatial and temporal pattern of mammalian auditory hair cell differentiation downstream of hedgehog signaling. J. Neurosci. 34, 12865–12876 (2014).

43. Doetzlhofer, A. et al. Hey2 regulation by FGF provides a Notch-independent mechanism for maintaining pillar cell fate in the Organ of Corti. Dev. Cell 16, 58–69 (2009).

44. Kamaid, A., Neves, J. & Giraldez, F. Id gene regulation and function in the prosensory domains of the chicken inner ear: A link between Bmp signaling and Atoh1. J. Neurosci. 30, 11426–11434 (2010).

45. Yu, R., Wang, P. & Chen, X.-W. The role of gfi1.2 in the development of zebrafish inner ear. Hear. Res. 396, 108055 (2020).

46. Jones, J. M. Inhibitors of differentiation and DNA binding (Ids) regulate Math1 and hair cell formation during the development of the Organ of Corti. J. Neurosci. 26, 550–558 (2006).

47. Hertzano, R. et al. Lhx3, a LIM domain transcription factor, is regulated by Pou4f3 in the auditory but not in the vestibular system. Eur. J. Neurosci. 25, 999–1005 (2007).

48. Hertzano, R. et al. Transcription profiling of inner ears from Pou4f3 ddl/ddl identifies Gfi1 as a target of the Pou4f3 deafness gene. Hum. Mol. Genet. 13, 2143–2153 (2004).

49. Deng, M. et al. LMO4 functions as a negative regulator of sensory organ formation in the mammalian cochlea. J. Neurosci. 34, 10072–10077 (2014).

50. Bae, S., Bessho, Y., Hojo, M. & Kageyama, R. The bHLH gene Hes6, an inhibitor of Hes1, promotes neuronal differentiation. Development 127, 2933–2943 (2000).

51. Fior, R. & Henrique, D. A novel hes5/hes6 circuitry of negative regulation controls Notch activity during neurogenesis. Dev. Biol. 281, 318–333 (2005).

52. Matern, M. S. et al. GFI1 functions to repress neuronal gene expression in the developing inner ear hair cells. Development 147, dev186015 (2020).

53. Hou, K. et al. A critical E-box in Barhl1 3′ enhancer is essential for auditory hair cell differentiation. Cells 8, 458 (2019).

54. Chonko, K. T. et al. Atoh1 directs hair cell differentiation and survival in the late embryonic mouse inner ear. Dev. Biol. 381, 401–410 (2013).

55. Waldhaus, J. et al. Stemness of the Organ of Corti relates to the epigenetic status of Sox2 enhancers. PLoS ONE 7, e36066 (2012).

56. Booth, K. T. et al. Novel loss-of-function mutations in COCH cause autosomal recessive nonsyndromic hearing loss. Hum. Genet. 139, 1565–1574 (2020).

57. Aibar, S. et al. SCENIC: single-cell regulatory network inference and clustering. Nat. Methods 14, 1083–1086 (2017).

58. Marchal, L., Luxardi, G., Thomé, V. & Kodjabachian, L. BMP inhibition initiates neural induction via FGF signaling and Zic genes. Proc. Natl. Acad. Sci. 106, 17437–17442 (2009).

59. Bienvenu, F. et al. Transcriptional role of cyclin D1 in development revealed by a genetic–proteomic screen. Nature 463, 374–378 (2010).

60. Luo, Z., Zhang, J., Qiao, L., Lu, F. & Liu, Z. Mapping genome-wide binding sites of Prox1 in mouse cochlea using the CUT&RUN approach. Neurosci. Bull. 37, 1703–1707 (2021).

61. Popova, E. Y. et al. Developmentally regulated linker histone H1c promotes heterochromatin condensation and mediates structural integrity of rod photoreceptors in mouse retina. J. Biol. Chem. 288, 17895–17907 (2013).

62. Freeman, S. D. & Daudet, N. Artificial induction of Sox21 regulates sensory cell formation in the embryonic chicken inner ear. PLoS ONE 7, e46387 (2012).

63. Mali, R. S. et al. FIZ1 is part of the regulatory protein complex on active photoreceptor-specific gene promoters in vivo. BMC Mol. Biol. 9, 87 (2008).

64. Pratapa, A., Amogh Jalihal, Law, J., Bharadwaj, A., & T M Murali. Benchmarking algorithms for gene regulatory network inference from single-cell transcriptomic data. (2020) doi:10.5281/ZENODO.3701939.

65. The ENCODE Project Consortium. An integrated encyclopedia of DNA elements in the human genome. Nature 489, 57–74 (2012).

66. Oki, S. et al. ChIP-Atlas: A data-mining suite powered by full integration of public ChIP-seq data. EMBO Rep. 19, (2018).

67. Xu, H., Ang, Y.-S., Sevilla, A., Lemischka, I. R. & Ma’ayan, A. Construction and validation of a regulatory network for pluripotency and self-renewal of mouse embryonic stem cells. PLoS Comput. Biol. 10, e1003777 (2014).

68. Simonyan, K. & Zisserman, A. Very deep convolutional networks for large-scale image recognition. in (2015). doi:arXiv:1409.1556v6.

69. Kuleshov, M. V. et al. Enrichr: a comprehensive gene set enrichment analysis web server 2016 update. Nucleic Acids Res. 44, W90–W97 (2016).

70. Blum, M. et al. The InterPro protein families and domains database: 20 years on. Nucleic Acids Res. 49, D344–D354 (2021).

71. Stuart, T. et al. Comprehensive integration of single-cell data. Cell 177, 1888–1902.e21 (2019).

72. Fang, R. et al. Comprehensive analysis of single cell ATAC-seq data with SnapATAC. Nat. Commun. 12, 1337 (2021).

73. Navarro Gonzalez, J. et al. The UCSC Genome Browser database: 2021 update. Nucleic Acids Res. 49, D1046–D1057 (2021).

74. Bailey, T. L. STREME: accurate and versatile sequence motif discovery. Bioinformatics 37, 2834–2840 (2021).

75. Khan, A. et al. JASPAR 2018: update of the open-access database of transcription factor binding profiles and its web framework. Nucleic Acids Res. 46, D260–D266 (2018).

76. Hume, M. A., Barrera, L. A., Gisselbrecht, S. S. & Bulyk, M. L. UniPROBE, update 2015: new tools and content for the online database of protein-binding microarray data on protein–DNA interactions. Nucleic Acids Res. 43, D117–D122 (2015).

77. Jolma, A. et al. DNA-binding specificities of human transcription factors. Cell 152, 327– 339 (2013).

78. Gupta, S., Stamatoyannopoulos, J. A., Bailey, T. L. & Noble, W. Quantifying similarity between motifs. Genome Biol. 8, R24 (2007).

